# RESTORING CONNEXIN-36 FUNCTION IN DIABETOGENIC ENVIRONMENTS PRECLUDES MOUSE AND HUMAN ISLET DYSFUNCTION

**DOI:** 10.1101/2020.11.03.366179

**Authors:** Joshua R St. Clair, Matthew J Westacott, Jose Miranda, Nikki L Farnsworth, Vira Kravets, Wolfgang E Schleicher, JaeAnn M Dwulet, Claire H Levitt, Audrey Heintz, Nurin WF Ludin, Richard KP Benninger

## Abstract

The secretion of insulin from β-cells in the islet of Langerhans is governed by a series of metabolic and electrical events, which can fail during the progression of type2 diabetes (T2D). β-cells are electrically coupled via Cx36 gap junction channels, which coordinates the pulsatile dynamics of [Ca^2+^] and insulin release across the islet. Factors such as pro-inflammatory cytokines and free fatty acids disrupt gap junction coupling under invitro conditions. Here we test whether gap junction coupling and coordinated [Ca^2+^] dynamics are disrupted in T2D, and whether recovery of gap junction coupling can recover islet function. We examine islets from donors with T2D, from db/db mice, and islets treated with proinflammatory cytokines (TNF-α, IL-1β, IFN-ɣ) or free fatty acids (palmitate). We modulate gap junction coupling using Cx36 over-expression or pharmacological activation via modafinil. We also develop a peptide mimetic (S293) of the c-terminal regulatory site of Cx36 designed to compete against its phosphorylation. Cx36 gap junction permeability and [Ca^2+^] dynamics were disrupted in islets from both human donors with T2D and db/db mice, and in islets treated with proinflammatory cytokines or palmitate. Cx36 over-expression, modafinil treatment and S293 peptide all enhanced Cx36 gap junction coupling and protected against declines in coordinated [Ca^2+^] dynamics. Cx36 over-expression and S293 peptide also reduced apoptosis induced by proinflammatory cytokines. Critically S293 peptide rescued gap junction coupling and [Ca^2+^] dynamics in islets from both db/db mice and a sub-set of T2D donors. Thus, recovering or enhancing Cx36 gap junction coupling can improve islet function in diabetes.

**KEY POINTS:**

- Cx36 gap junction permeability and associated coordination of [Ca2+] dynamics is diminished in human type 2 diabetes (T2D) and mouse models of T2D.
- Enhancing Cx36 gap junction permeability protects against disruptions to the coordination of [Ca2+] dynamics.
- A novel peptide mimetic of the Cx36 c-terminal regulatory region protects against declines in Cx36 gap junction permeability.
- Pharmacological elevation in Cx36 or Cx36 peptide mimetic recovers [Ca2+] dynamics and GSIS in human T2D and mouse models of T2D.

## INTRODUCTION

Type2 diabetes (T2D) results from β-cell failure in the context of obesity-related insulin resistance, such that insulin production does not meet physiological demands. Many patients with T2D require β-cell secretagogues or exogenous insulin therapy (Turner *et al*., 1999; van Raalte & Diamant, 2011) following the loss of first phase insulin secretion and reduced second phase insulin dynamics (Meier *et al*., 2005; Menge *et al*., 2011). While the etiology of T2D is multifactorial, it is becoming increasingly clear that T2D develops in patients after the onset of β-cell dysfunction (Pfeifer *et al*., 1981; Turner *et al*., 1999; Prentki & Nolan, 2006; Ray *et al*., 2009; van Raalte & Diamant, 2011). Both reduced β-cell mass as well as intrinsic β-cell dysfunction (de Koning *et al*., 2008; Bonner-Weir & O’Brien, 2008) have been shown to exist prior to clinical onset of disease (Xu *et al*., 2003; Abdulreda & Berggren, 2013). Therefore, therapeutics that maintain β-cell mass and function during T2D progression have the potential to prevent disease development. While inflammation and lipotoxicity are implicated in T2D, the mechanisms that determine the advancement of islet dysfunction in pre-T2D are poorly understood.

Pancreatic islets of Langerhans maintain blood glucose homeostasis by producing and secreting insulin. Insulin secretion is triggered by glucose-stimulated increases in intracellular free-calcium ([Ca^2+^]) (Prentki *et al*., 1997; Schuit *et al*., 2001; Benninger & Piston, 2014). Glucose-stimulated [Ca^2+^] increases are in the form of oscillations that generate pulses of insulin release, which are important for proper insulin action and glucose utilization (Meier *et al*., 2005; Bertram *et al*., 2007). Connexin-36 (Cx36) forms gap junctions (GJs) that electrically couple β-cells within islets (Ravier *et al*., 2005; Benninger *et al*., 2008). Loss-of-function studies have demonstrated that Cx36 GJs are critical for controlling oscillatory [Ca^2+^] across the islet to generate pulsatile insulin secretion under elevated glucose (Ravier *et al*., 2005; Benninger *et al*., 2008; Head *et al*., 2012), as well as to dampen spontaneous β-cell electrical activity and inhibit insulin secretion under low glucose (Speier *et al*., 2007; Benninger *et al*., 2011). In the absence of Cx36, there is a complete absence of gap junction electrical coupling, and disorganized and unsynchronized [Ca^2+^] oscillations lead to a complete loss of first phase insulin secretion, diminished second phase pulses and glucose intolerance (Ravier *et al*., 2005; Head *et al*., 2012). Furthermore, proper GJ coupling has been shown to be important for protecting against β-cell apoptosis and maintaining β-cell mass (Klee *et al*., 2011b, 2011a). While a direct role for GJ coupling in T2D progression remains unclear, the physiological consequences of islet uncoupling by a genetic disruption of Cx36 is analogous to pathological observations of patients T2D (Pfeifer *et al*., 1981; Menge *et al*., 2011), including glucose intolerance and altered insulin secretion dynamics suggesting a role for islet GJ uncoupling in the disease.

Chronic inflammation and gluco-lipotoxicity as a result of insulin resistance and metabolic stress in T2D has detrimental effects on β-cell function, including dysregulated [Ca^2+^] dynamics, reduced insulin synthesis and secretion, and increased β-cell apoptosis (Lee *et al*., 1994; McGarry, 2002; El-Assaad *et al*., 2003; Boden, 2005, 2008; Poitout *et al*., 2006; Hatanaka *et al*., 2014). For example, in obese individuals, excessive adipose mass releases increasing amounts of FFAs and pro-inflammatory cytokines into the bloodstream (Xu *et al*., 2003; Donath *et al*., 2008; Imai *et al*., 2016). These pro-inflammatory cytokines include tumor necrosis factor alpha (TNF-α) (Hotamisligil *et al*., 1993), interleukin (IL)-6 (Mahdi *et al*., 2012), IL-1β (Maedler *et al*., 2017), among others (Tataranni & Ortega, 2005; Marchetti, 2016). Pro-inflammatory cytokines can induce GJ uncoupling within mouse and human islets (Farnsworth *et al*., 2016a). Chronic FFA exposure can also induce GJ uncoupling within mouse and human islets (Hodson *et al*., 2013). In animal models of obesity, dysregulated [Ca^2+^] dynamics have also been observed (Ravier *et al*., 2002; Corezola do Amaral *et al*., 2020). Taken together, this suggests a role for GJ uncoupling in the development of islet dysfunction in T2D. However, little is known about the impact of chronic FFA and inflammation on Cx36 GJ uncoupling and [Ca^2+^] in prior to T2D and whether in fact this uncoupling significantly impacts the progression of islet dysfunction.

Cx36 gap junction coupling protects β-cells from the susceptibility of apoptosis (Klee *et al*., 2011a). Together with the role for GJ coupling in regulating insulin secretion, maintenance of Cx36 function in diabetic islets may have a therapeutic benefit. Modulation of GJ function with structural mimetic peptides has been demonstrated in treatment of cardiac arrhythmias (Eloff *et al*., 2003), myocardial infarction and ischemia (Hawat *et al*., 2012). The C-terminus of Cx36 contains many sites that regulate GJ function, including phosphorylation by PKA or CaMKII (Urschel *et al*., 2006; Kothmann *et al*., 2007; Alev *et al*., 2008), or interactions with microtubules (Brown *et al*., 2019) and tight junctions (Flores *et al*., 2008). Both PKA and PKCδ has been demonstrated to regulate Cx36 GJ coupling within the islet (Hodson *et al*., 2013; Farnsworth *et al*., 2016a). Thus, we hypothesized that maintaining Cx36 GJ coupling during inflammatory and lipotoxic events in T2D may provide a novel mechanism by which insulin secretion and β-cell mass can be maintained during pre-diabetes, and thus delay or prevent disease onset.

In this study, we sought to determine the pro-inflammatory and FFA environmental factors that influence GJ and Cx36 function, and to identify a mechanism by which GJ uncoupling could be prevented in the face of diabetogenic factors. We show that GJ coupling, insulin secretion, and cell viability are all negatively affected by diabetogenic environments, including in human T2D, animal models of diabetes and by pro-inflammatory cytokines and FFAs. We further demonstrate that reducing or preventing the decline in Cx36 gap junction coupling by either pharmacological activators or by a novel mimetic inhibitory peptide can preclude this dysfunction.

## METHODS

### Animals

All experimental procedures were performed at the University of Colorado Anschutz Medical Campus, and were approved by the Institutional Care and Use Committee (IACUC). Commercially-available wildtype female C57BL/6NHsd (Envigo stock #044) were used aged 12-16 weeks. Diabetes prone C57BKS.Cg-Leprdb (“db/db”) mice were used aged 4-6 weeks (Jackson Laboratories stock #000642) with appropriate age- and sex-matched strain C57BLKS control mice (Jackson Laboratories stock #000662).

### Islet isolation and culture

For mouse islet isolation, animals were injected with ketamine/xylazine at 83/17 mg/kg to induce general anesthesia, and islets were isolated by a method previously described elsewhere following collagenase infusion, pancreas dissection and digestion, and hand picking of islets (Koster *et al*., 2002; Farnsworth *et al*., 2016a). Isolated islets were cultured overnight in RPMI medium 1640 (Life Technologies) supplemented with 10% FBS, 11mM glucose, 100 units/ml of penicillin, and 100 μg/ml of streptomycin at 37 °C under humidified 5% CO2. Human islets were received from the Integrated Islet Distribution Program, obtained from either healthy donors or donors with type2 diabetes (Table S1) and were cultured overnight in CMRL medium 1066 (Fisher Scientific) prior to initiating experiments.

### Islet treatments

For experiments involving treatment with pro-inflammatory cytokines, isolated mouse and human islets were treated for 1 hour in imaging buffer (125mM NaCl, 5.7mM KCl, 2.5mM CaCl2, 1.2mM MgCl2, 10mM HEPES, 0.1% bovine serum albumin (BSA), and 2mM glucose) or for 24 hours in either RPMI (mouse islets) or CMRL (human islets) based media. 1 hour treatments were used for imaging based measurements (FRAP for gap junction permeability, [Ca^2+^]), and 24 hour treatments used for viability and glucose-stimulated insulin secretion (GSIS) measurements. A “type-1” like cytokine cocktail consisted of 10 ng/ml of recombinant mouse tumor necrosis factor-α (TNF-α, R&D Systems, Minneapolis, MN), 5 ng/ml of recombinant mouse interleukin-1β (IL-1β, R&D Systems), and 100 ng/ml of recombinant mouse interferon-γ (IFN-γ, R&D Systems) (Farnsworth *et al*., 2016a). Where noted, a “type 2” like cytokine cocktail consisted of either IL1β+IL6 (1 ng/ml and 10 pg/ml respectively) or TNF-α+IL-1β (0.5 ng/ml and 1 ng/ml), which were used to mimic a proinflammatory cytokine environment in pre-T2 diabetes. For experiments involving treatment with free-fatty acid isolated mouse and human islets were incubated in 200 μM palmitate (bound to 0.17mM BSA) in either RPMI (mouse islets) or CMRL (human islets) for 24 hours or 48 hours. For experiments with an “untreated” condition, islets were maintained in RPMI/CMRL culture media prior to experiments, with no exchange of media.

### Fluorescence Recovery after Photobleaching

To assess β-cell gap junction permeability in mouse and human islets, fluorescence recovery after photobleaching (FRAP) was used as previously established and described (Farnsworth *et al*., 2014). Briefly, islets were immobilized on MatTek dishes (MatTek Corp., Ashland, MA) that were coated with CellTak (BD Biosciences, San Jose, CA) and stained with 12.5μm Rhodamine123 (Sigma) for 30 min at 37 °C. Islets were imaged either on a Zeiss LSM 510 Meta confocal microscope with a ×40 1.2NA water immersion objective, or a Zeiss LSM 800 with a x40 1.2NA water immersion objective. Rhodamine123 was excited at 488 nm and images were collected with a 488-nm narrow band-pass dichroic beam splitter, 490-nm long pass dichroic beam splitter, and a 505-nm long pass emission filter. Procedurally, half of the islet area was selected and photobleached for 235.5 s at 316.05 milliwatt/cm^2^ and the fluorescence recovery was measured in the bleached area. Recovery rates were calculated from the inverse exponential fluorescence recovery curve for the entire bleached area (Farnsworth *et al*., 2014). Previous work from our lab has demonstrated that this method measures GJ permeability in β-cells only, as GJs do not couple α-cells.

FRAP data was either presented as raw recovery rates calculated from the recovery curve, or as the recovery rate normalized to paired untreated islets from experiments performed from on the same day (where indicated).

### Intracellular Ca^2+^ Imaging and Analysis

Intra-cellular fee-calcium ([Ca^2+^]) dynamics were assessed in islets stained with 4μm Fluo-4 AM for 1 hour at 35 ± 1°C in imaging buffer (125mM NaCl, 5.7mM KCl, 2.5mM CaCl2, 1.2mM MgCl2, 10mM HEPES, 0.1% bovine serum albumin (BSA), and 2 mM glucose). Islets were imaged on MatTek dishes and were maintained at 37 °C with an Eclipse-Ti wide field microscope (Nikon) and a ×20 0.75 NA Plan Apo objective. Images were acquired at 1 frame per second five minutes after stimulation with 11 mm glucose, for 300 seconds. Fluo-4 was imaged using a 490/40-nm band-pass excitation filter and a 525/36-nm band-pass emission filter (Chroma).

The proportion of the islet that showed synchronized oscillations was determined over the 300s intervals, as previously described (Hraha *et al*., 2014; Westacott *et al*., 2017). For mouse islets, a reference time-course was taken averaged over the whole islet or sub-region of the islet. A cross-correlation coefficient was then defined between this reference and the time-course corresponding to each pixel within the image that showed significant Fluo4 staining over background. A region was defined as synchronized if the cross-correlation coefficient was >0.75. The area of the islet showing synchronized oscillations was normalized to the total are of the islet. For human islets, elevations in [Ca^2+^] were identified for time-course corresponding to each pixel within the image that showed significant Fluo4 staining over background. Contagious pixels that showed coincident elevations in [Ca^2+^] for at least 50% of the oscillations were defined as synchronized regions. The area of the largest synchronized region was normalized to the total are of the islet.

[Ca^2+^] data was either presented as % synchronized area, or as the synchronized area normalized to paired untreated islets from experiments performed from on the same day (where indicated).

### Islet Viability

Following 24-h treatment in indicated condition, islets were incubated with a 1:1000 dilution of LIVE/DEAD™ Fixable Near-IR Dead Cell Stain (Life Technologies) diluted in HBSS for 30 min at room temperature. Islets were imaged on a Zeiss LSM 800 Meta confocal microscope with a ×40 1.2NA water immersion objective. The dye was excited with a 633-nm He-Ne laser with a UV/488/543/633-nm dichroic beam splitter, and images for live/dead cells were collected with a 650-710nm band-pass emission filter. Viability was calculated from manual counts of live and dead cells averaged over 3 images of increasing depth in each islet.

### Glucose stimulated insulin secretion assay (GSIS)

Glucose stimulated insulin secretion (GSIS) was determined in static assays, as described previously (Farnsworth *et al*., 2016a; Westacott *et al*., 2017). Following 24-hour of experimental treatments, duplicates of ∼5 IEQs per tube (mouse islets) and ∼10 IEQs (human islets) were incubated in 500μl of Krebs-Ringer buffer (128.8mM NaCl, 5mM KCl, 1.2mM KH2PO4, 2.5mM CaCl2, 1.2mM MgSO4, 1mM HEPES, 0.1% BSA, pH 7.4) with 2mM glucose for 1 hour, followed by 1 hour stimulation with either 2mM or 20mM glucose. Islets were lysed by 2% Triton X-100 in deionized water followed by a freeze-thaw cycle. Both supernatant and islet lysate were collected for analysis of insulin secretion and content, respectively. Secreted insulin concentrations from both mouse and human GSIS assays were measured with a mouse ultrasensitive insulin ELISA kit (ALPCO; cross-reactive for human insulin) per the manufacturer’s instructions. GSIS data are reported as % insulin secretion (supernatant) normalized to insulin content (lysate), or insulin secretion at 20mM glucose normalized to insulin secretion at 2mM glucose (where indicated).

### Cx36 Mimetic Peptide Design

An eleven amino acid sequence corresponding to residues 288-298, flanking residue 293 (serine) of murine connexin-36 (QAKRK<u>S</u>VYEIR), was generated to mimic this portion of the c-terminus (“S293”). A 16-amino acid antennapedia domain (RQIKIWFQNRRMKWKK) was included on the N-terminus, immediately preceding the connexin-36 11 amino acid sequence to facilitate cellular internalization <u>(Avignolo *et al*., 2008)</u>. A biotin tag was included on the N-terminus, and amide domain on the c-terminus, as shown in Figure 4A. This peptide was synthesized by the University of Colorado Peptide and Protein Chemistry Core Facility (shown in Figure 4A). A scrambled peptide (“Scr-S293”) was synthesized as a negative control (sequence: KYQREKVARIS). The sequence was manually designed by the CU-PPCC faculty. Scrambled peptide control experiments were performed alongside all S293 experiments to ensure peptide specificity and to account for off-target effects of non-Cx36 peptide sequences.

### Statistical Analysis

All data presented represent the average over all mice for each measurement. Unless indicated, error bars represent either the mean ± s.e.m (with symbols representing individual datapoints) or mean ± S.D. For human islets, data is presented as individual points per donor, together with the mean ± s.e.m. Statistical significance was determined using either a Student’s t-test or an analysis of variance (ANOVA) with Tukey’s post hoc analysis. An α of 0.05 was pre-determined prior to experiments to be statistically significant, and statistical significance is indicated where appropriate.

## RESULTS

### Reduced GJ function and [Ca^2+^] coordination in human islets from T2D donors

To determine the role of β-cell gap junction (GJ) coupling in the pathogenesis of type2 diabetes (T2D), we first assessed GJ permeability and [Ca^2+^] dynamics in human islets from confirmed T2D donors and compared these with islets from age- and BMI-matched non-diabetic donors. Islets from T2D human donors exhibited a significant 28% reduction in recovery rate (reflecting GJ permeability, (Farnsworth *et al*., 2014)) compared to islets from non-diabetic human donors (Figure 1A, p=0.0012). There was no correlation between GJ permeability and HbA1c among the T2D group (Figure S1), indicating the decrease in coupling was unlikely to result from glucose toxicity but is caused by some other factor related to T2D progression.

**Figure 1:**
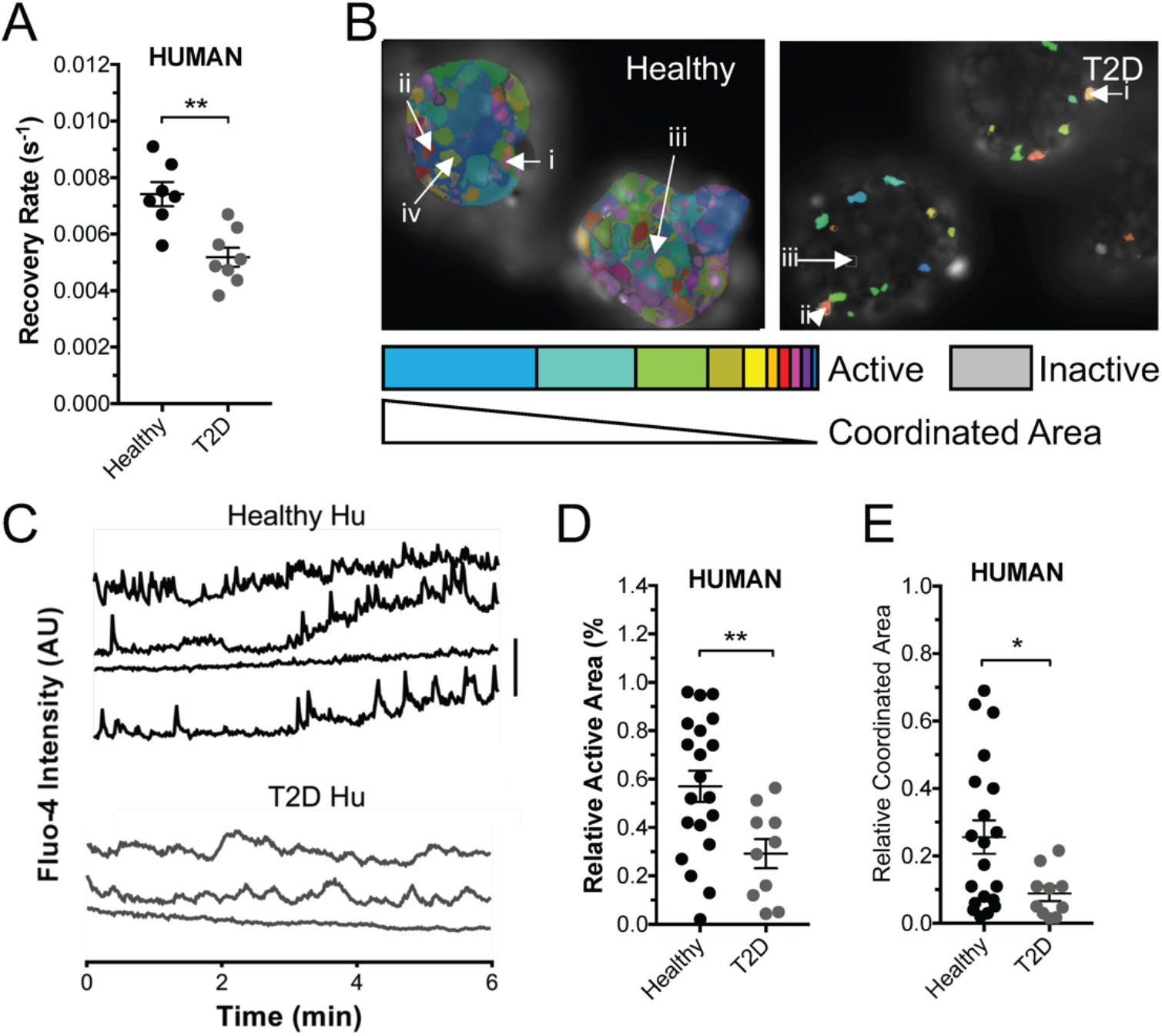
Reduced GJ function and [Ca2+] coordination in human islets from T2D donors. A) Gap junction permeability, as measured by FRAP recovery rate in islets from T2D donors and from age and BMI matched healthy donors. B) Representative false color map of [Ca^2+^] activity and coordination in human islet from a healthy donor (top) and from a T2D donor (bottom). [Ca^2+^] activity is represented by presence of false color, with each color representing a separate region of [Ca^2+^] coordination, as indicated in legend below. C) Time courses from individual cells (i–iv or i-iii) indicated in each of these islets. D) Area of [Ca^2+^] activity normalized to islet size (relative active area) averaged over human islets from each healthy and T2D donor. E) Largest area of coordinated [Ca^2+^] activity normalized to islet size (relative coordinated area) averaged over human islets from each healthy and T2D donor. Data in A representative of n=7 healthy donors and n=8 T2D donors. Data in D,E representative of n=20 healthy donors and n=10 T2D donors. In A ** represents p=0.0012, in D ** represents p=0.0099, in E * represents p=0.029, comparing groups indicated (all unpaired Student’s t-test). Data in A,D,E presented as one point per donor.

GJ coupling coordinates oscillatory [Ca^2+^] dynamics across the islet. We quantified [Ca^2+^] dynamics in T2D human islets and compared them those dynamics measured from age- and BMI-matched non-diabetic controls (Figure 1B,C). Islets from T2D human donors had significantly reduced proportion of cells that showed elevated [Ca^2+^] (Figure 1D, ‘active area’: p=0.0099). Islets from T2D human donors also had a significantly reduced proportion of cells that showed synchronized [Ca^2+^] dynamics (Figure 1E, ‘coordinated area’: p=0.029). Again, there was no correlation between measures of [Ca^2+^] dynamics with HbA1c among the T2D donor islet group (Figure S1). We also did not observe a correlation between fold-change in glucose-stimulated insulin secretion (GSIS) under static assay, with HbA1c (Figure S1). GSIS was similar between healthy and T2D groups (healthy 2.34±0.24, T2D 1.96±0.36, p=0.36). Taken together, these data indicate, for the first time, that islets from confirmed T2D humans have reduced gap junction function, which logically leads to disrupted coordinated [Ca^2+^] dynamics.

### Diabetogenic environments disrupt GJ function and [Ca^2+^] coordination

Given results from human T2D islets, we sought to understand the pathophysiological mechanisms for reduced GJ coupling. To this end, we performed similar experiments with islets isolated from the db/db mouse model of T2D: an animal model of obesity induced diabetes that results from leptin signaling deficiency. We examined db/db mice at 4 weeks of age, where animals were euglycemic, but glucose intolerant, and thus represent a “pre-diabetic” phase prior to marked β-cell decline. Compared to control animals (C57BLKS), islets from 4 week old db/db mice had significant reduction in GJ function as assessed with FRAP (Figure 2A; p=0.049). db/db islets also exhibited disorganized [Ca^2+^] elevations (Figure 2B), and reduced [Ca^2+^] synchronization compared to control islets (Figure 2C; p=0.002). Together with data from T2D human islets, the FRAP and [Ca^2+^] data from db/db mouse islets indicates GJ uncoupling as a common pathophysiological feature in both human and mouse T2D.

**Figure 2:**
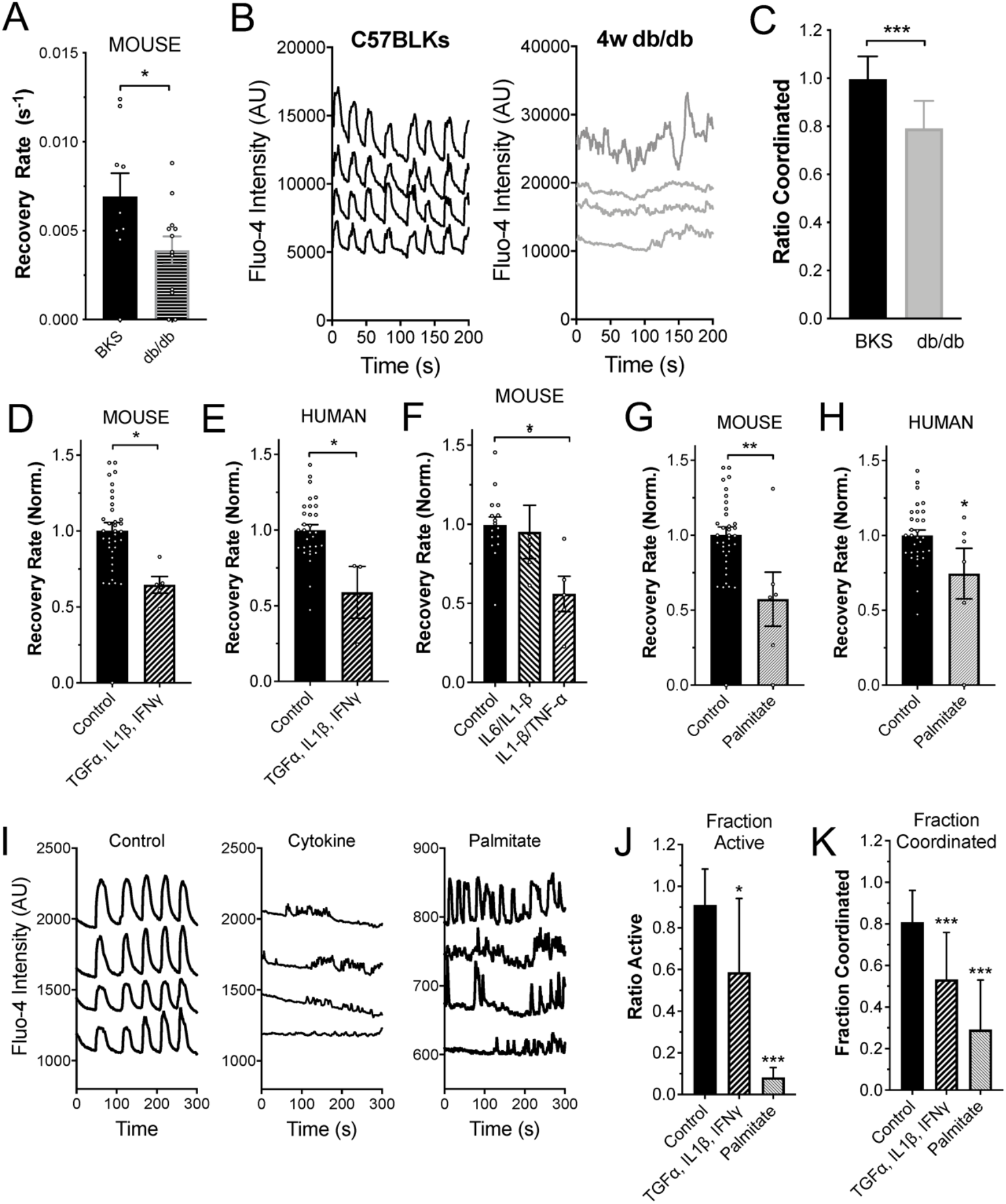
Diabetogenic environments disrupt GJ function and [Ca^2+^] coordination. A) Mean gap junction permeability, as measured by FRAP recovery rate, in islets from 4w C57BLKS (BKS) controls and 4w db/db mice. B) Representative [Ca^2+^] time courses from four individual cells within an islet from a 4w C57BLKS control and 4w db/db mouse. C) Mean proportion of islets that show synchronized Ca^2+^ oscillations, normalized to the mean value observed in islets from BKS mice measured on the same day. D) Mean gap junction permeability, in untreated islets and islets treated with a cocktail of 1 ng/ml mouse recombinant TNF-α, 0.5 ng/ml IL-1β, 10 ng/ml IFN-γ. Data is normalized to the paired untreated islets. E) As in D for human islets treated with the same cytokine cocktail of TNF-α, IL-1β, IFN-γ. F) As in D for mouse islets treated with a cocktail of 1 ng/ml mouse recombinant TNF-α, 0.5 ng/ml IL-1β, or with a cocktail of 1 ng/ml IL1β, 10 pg/ml IL6. G) Mean gap junction permeability, in untreated mouse islets and islets treated with 200μM palmitate (bound to 0.17mM BSA). Data is normalized to the paired untreated islets. H) As in G for human islets treated with palmitate. I) Representative [Ca^2+^] time courses from four individual cells within an untreated islet (left), for an islet treated with TNF-α, IL-1β, IFN-γ (middle, as in D) or for an islet treated with palmitate (right, as in G). J) Area of [Ca^2+^] activity normalized to islet size (fraction active). K) Area of coordinated [Ca^2+^] normalized to islet size (fraction coordinated). Data in A representative of n=9 C57BLKS and n=12 db/db islets. Data in C representative of n=24 C57BLKS and n=5 db/db islets. Data in D representative of n=10 untreated and n=6 cytokine treated islets. Data in E representative of n=7 untreated and n=6 cytokine treated islets. Data in F representative of n=16 untreated and n=5 cytokine treated islets. Data in G representative of n=6 untreated and n=5 palmitate treated islets. Data in H representative of n=6 untreated and n=6 palmitate treated islets. In A * represents p=0.049; in C *** represents p=0.002; In D* represents p=0.020; in E * represents p=0.0066; in F * represents p=0.047; in G * represents p=0.007; in H * represents p=0.025; in J * represents p=0.0012, *** represents p<0.0001; in K *** represents p=0.0003 and p=<0.0001, comparing groups indicated (all unpaired Student’s t-test). Data in A, D-H presented as mean±s.e.m. with each data point representing an islet. Data in C,J,K presented as mean±SD.

To delineate the diabetogenic mechanisms that lead to GJ uncoupling in T2D donor human and db/db mouse islets, we mimicked a diabetogenic environment by treating wildtype mouse islets and human islets from healthy donors with several cocktails of pro-inflammatory cytokines, as well as with palmitate, a long-chain free fatty acid. A cytokine cocktail of TNF-α, IL-1β, and IFN-γ reduced islet GJ uncoupling in both mouse and healthy human islets to a similar degree when normalized to untreated islets, as we have previously observed (Figure 2D, p=0.020; Figure 2E, p=0.0066). A cytokine cocktail of IL-1β, and TNF-α alone also reduced GJ coupling when normalized to untreated islets (Figure 2F, p=0.047). However, a cytokine cocktail of IL-1β and IL6 did not induce significant GJ uncoupling (Figure 2F). Finally, we assessed the ability of the long-chain free fatty acid palmitate to induce changes in GJ coupling. Treatment with 200 μM palmitate resulted in significantly reduced GJ coupling in both mouse islets (Figure 2G, 0.57±0.18, p=0.007) and human islets (Figure 2H, 0.74±0.17, p=0.025). These data clearly suggest that both IL-1β/TNF-α and/or free fatty acids have effects on mediating GJ uncoupling in the pre-T2D clinical milieu. These treatments that reduced GJ coupling also led to reductions in [Ca^2+^] activity and [Ca^2+^] coordination in mouse pancreatic islets treated in the same manner (Figure 2I-K).

Thus, convergent mechanisms may exist within the diabetogenic environment, such that both pro-inflammatory cytokine cocktails and FFAs (palmitate) induce GJ uncoupling and [Ca^2+^] desynchronization.

### Increasing Cx36 gap junction coupling protects against islet decline and cell death

Gap junctions in both mouse and human islets are formed by Cx36. To test whether elevating Cx36 gap junction coupling protects against islet decline under diabetogenic conditions we first over-expressed Cx36 in β-cells via a transgenic mouse, as previously characterized (Klee *et al*., 2011a). In islets from over-expression mice, Cx36 gap junction coupling was elevated compared to islets from age matched or littermate wild-type controls (Figure 3A). Following treatment with the cytokine cocktail of TNF-α, IL-1β, and IFN-γ Cx36 GJ coupling was elevated compared to controls (Figure 3A). Cell death was significantly less in islets from Rip-Cx36 mice compared to control mice, following treatment with the cytokine cocktail (Figure 3B), as previously observed (Klee *et al*., 2011a). While islets from Rip-Cx36 mice showed a trend to elevated GSIS compared to islets from non-transgenic controls, this increase was not statistically significant (Figure 3C).

**Figure 3:**
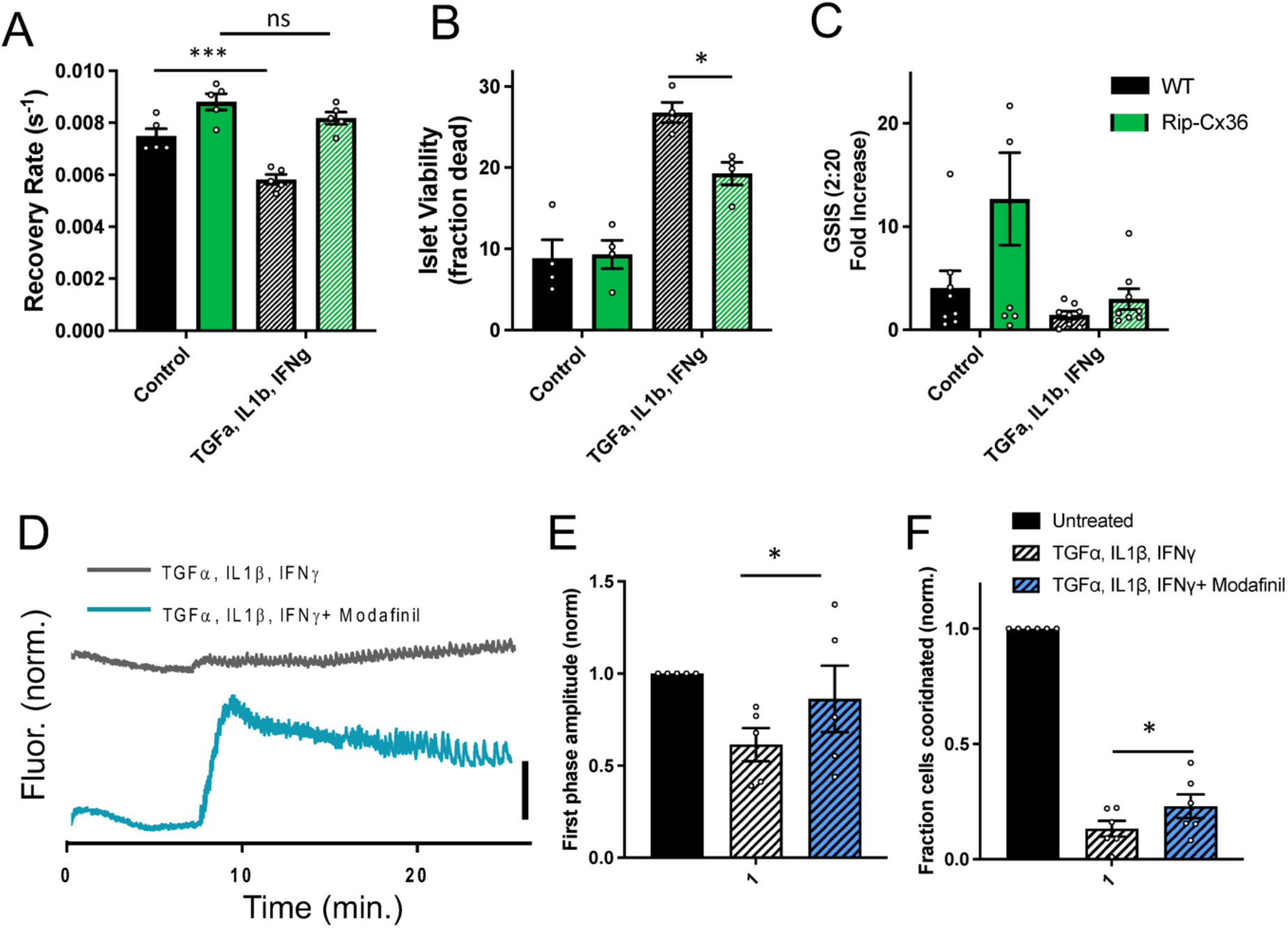
Increasing Cx36 gap junction coupling protects against islet dysfunction and cell death. A) Mean gap junction permeability, as measured by FRAP recovery rate, in islets from wild-type and Rip-Cx36 overexpression mice, that are either untreated or treated with a cocktail of 1 ng/ml mouse recombinant TNF-α, 0.5 ng/ml IL-1β, 10 ng/ml IFN-γ. B) Islet viability, as indicated by the proportion of cells labelled as dead, in islets as in A. C) Glucose-stimulated Insulin secretion (GSIS) in islets as in A. D) Representative [Ca^2+^] time courses averaged over an islet treated with TNF-α, IL-1β, IFN-γ as in A-C (grey) or an islet treated with TNF-α, IL-1β, IFN-γ as in A-C plus 50 μM modafinil. E) Amplitude of [Ca^2+^] elevation ∼10 minutes after glucose elevation, for untreated islets and treatments in D. Data is normalized to the elevation measured in untreated islets. F) Area of coordinated [Ca^2+^] normalized to islet size (fraction coordinated), for untreated islets and treatments in D. Data is normalized to the elevation measured in untreated islets. Data in A representative of n=5 wild-type and n=5 Rip-Cx36 mice; data in B representative of n=4 wild-type and n=4 Rip-Cx36 mice; data in C representative of n=8 wild-type and n=8 Rip-Cx36 mice; data in E representative of n=5 mice for each treatment, data in F representative of n=6 mice for each treatment. In A * represents p=0.0007, ns represents p=0.21; in B * represents p=0.02; in E * represents p=0.04; in B * represents p=0.05; comparing groups indicated (all unpaired Student’s t-test). Data in A-C, E-F presented as mean±s.e.m. with each data point representing an islet.

Modafinil has been shown to elevate Cx36 GJ coupling in the islet (Farnsworth *et al*., 2014; Westacott *et al*., 2017), albeit with the mechanisms poorly understood (Urbano *et al*., 2007). Modafinil treatment significantly elevated both the first phase elevation in [Ca^2+^] (Figure 3D,E) and synchronization of Ca^2+^ oscillations (Figure 3D,F) following treatment with the cytokine cocktail. Therefore elevating Cx36 GJ coupling through genetic or pharmacological means provides protection against disruptions to [Ca^2+^] dynamics and blunts cell death induced by diabetogenic conditions.

### Cx36-S293 is critical for GJ uncoupling in diabetogenic environments

Cx36 GJ coupling can be regulated by phosphorylation of various residues on the protein, particularly on the intra-cellular loop and c-terminus (Urschel *et al*., 2006; Li *et al*., 2014; Ivanova *et al*., 2015; Zhang *et al*., 2015). Phosphorylation of Cx36 at serine-293 (S293) by PKA was shown to induce Cx36 uncoupling in AII Amacrine cells in the mouse retina, as determined by dye tracer experiments (Urschel *et al*., 2006; Kothmann *et al*., 2009). Furthermore, dynamic regulation of Cx35 coupling by phosphorylation and de-phosphorylation at serine-273 in the zebrafish (analogous to S293 on the mammalian Cx36) retina is regulated by PKA (Kothmann *et al*., 2007). Our lab has also demonstrated a role for PKCδ in regulating β-cell Cx36 GJ coupling (Farnsworth *et al*., 2016a). Given the role of PKA and PKCδ in regulating Cx36 (in the islet and elsewhere), we hypothesized that prevention of Cx36-S293 phosphorylation in the context of the diabetogenic environment described above could lead to preservation of GJ coupling and downstream factors such as [Ca^2+^] dynamics. To this end, we synthesized a novel mimetic inhibitory peptide targeted toward S293 (“S293”) on the Cx36 c-terminus (Figure 4A). This synthetic peptide included an antennapedia internalization domain (Avignolo *et al*., 2008); an 11-amino acid sequence corresponding to the Cx36 sequence flanking S293 (Cx36:288-298); and an N-terminal biotin tag. Effective intracellular internalization of the peptide occurred in dissociated β-cells from mouse islets (Figure 4B), as measured by fluorescence imaging of an Alexa-488 tagged streptavidin that bound to the biotin tag on the S293 peptide. We also synthesized a peptide with internalization domain and biotin tag, but with a scrambled sequence of Cx36:288-298 (“Scr-S293”).

**Figure 4:**
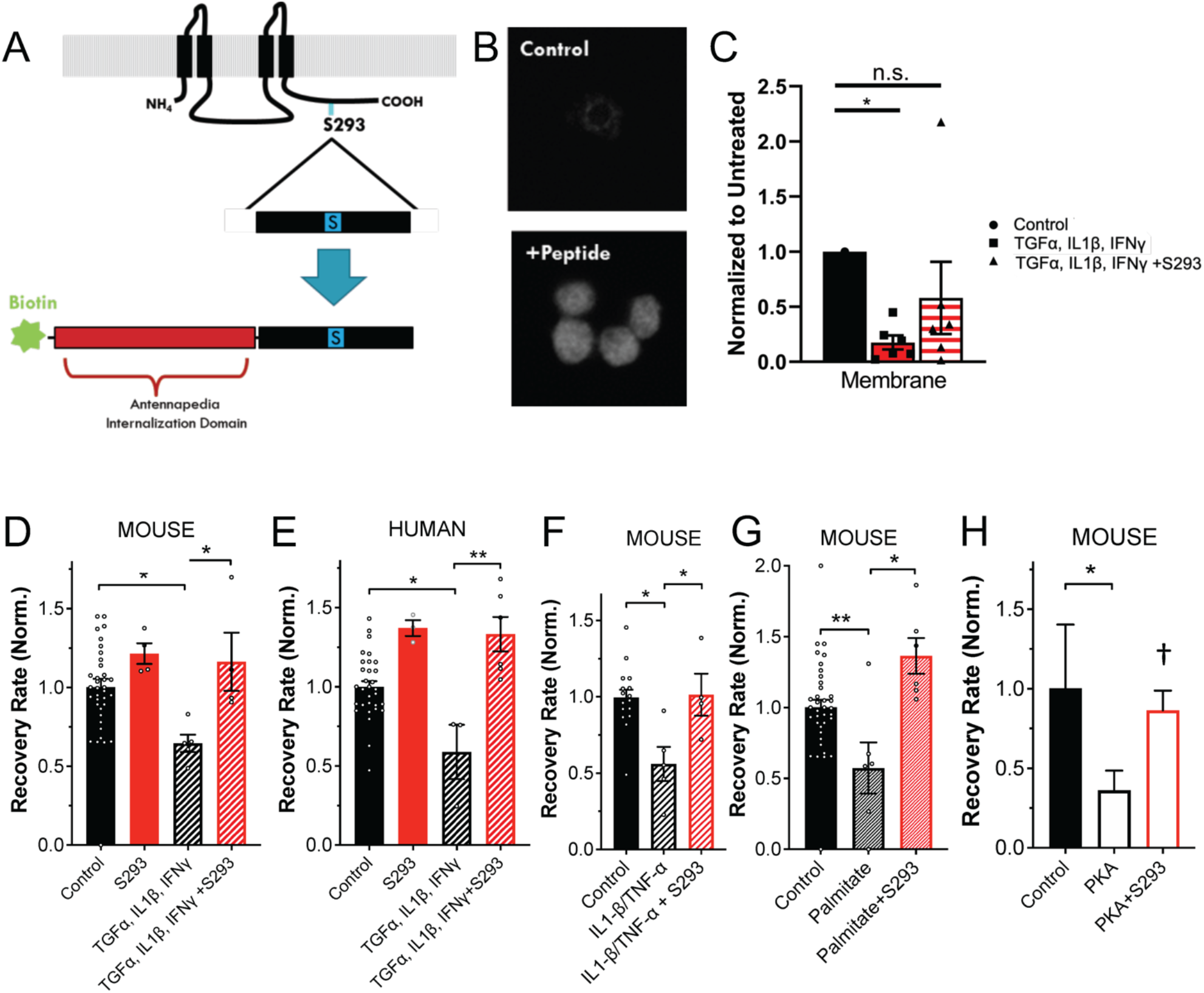
Cx36-S293 is critical for GJ uncoupling in diabetogenic environments. A) Schematic of S293 peptide mimetic of Cx36 c-terminus. B) Fluorescence image of biotin labelling in control, untreated cells and cells treated with S293 mimetic. C) Mean intensity of membrane-localized Cx36, quantified from western blot of membrane fraction of mouse islets treated with a cocktail of 1 ng/ml mouse recombinant TNF-α, 0.5 ng/ml IL-1β, 10 ng/ml IFN-γ and/or 1 μM S293 peptide. D) Mean gap junction permeability, as measured by FRAP recovery rate, in mouse islets treated with a cocktail of 1 ng/ml mouse recombinant TNF-α, 0.5 ng/ml IL-1β, 10 ng/ml IFN-γ and/or 1 μM S293 peptide. Data is normalized to the paired untreated islets. E) As in D for human islets. F) As in D for mouse islets treated with a cocktail of 1 ng/ml mouse recombinant TNF-α, 0.5 ng/ml IL-1β, and/or 1 μM S293 peptide. G) As in D for mouse islets treated with 200μM palmitate (bound to 0.17mM BSA) and/or 1 μM S293 peptide. H) As in D for mouse islets treated with 300uM 6-Bnz-cAMP (PKA) and/or 1 μM S293 peptide. Data in C representative of n=6 experiments; data in D representative of islets from n=5 mice; data in E representative of islets from n=6 donors; data in F representative of islets from n=4 mice; data in G representative of islets from n=7 mice; data in H representative of islets from n=2 mice. In C * represents p<0.01 D * represents p=0.021; in E ** represents p=0.007; in F * represents p=0.036; in G * represents p=0.004; in H * represents p=0.037, † represents p=0.056; comparing groups indicated (all unpaired Student’s t-test). Data in C-H presented as mean±s.e.m. with each data point representing an islet. Data in H presented as mean±SD.

We tested the function of the S293 peptide and whether it could block the GJ uncoupling induced by the cocktails of pro-inflammatory cytokines and palmitate (Figure 2), as well as the reduced membrane localization (Farnsworth *et al*., 2016b). Incubation of islets with the S293 peptide reduced the loss of membrane localized Cx36 (Figure 4C), and prevented the cytokine-mediated decreases in GJ permeability in response to TNF-α, IL-1β, and IFN-γ in mouse islets (Figure 4D, p= 0.021) and in human islets (Figure 4E, p=0.007). We observed a small increase in GJ permeability observed when human islets were treated with S293 peptide alone (Figure 4E, 37±4% increase, p=0.03), but not in the case of mouse islets (Figure 4D, 22±6% increase, p=0.33). The S293 inhibitory peptide was effective in preventing cytokine-mediated (TNF-α, IL-1β, and IFN-γ) decreases in GJ permeability, down to 30 nM concentration, highlighting its potency (Figure S2A). Similarly, incubation of mouse islets with S293 peptide in the presence of TNF-α, IL-1β alone prevented decreases in GJ permeability (Figure 4F, p=0.036).

We also tested whether the S293 peptide could prevent free-fatty acid (100μM palmitate) mediated decreases in GJ coupling. Decreases in GJ coupling were prevented in islets treated with palmitate and S293 peptide, when compared to islets treated with palmitate alone (Figure 4G, p=0.004). PKA activation has been suggested to mediate the decrease in GJ coupling induced by elevated free-fatty acids. Incubation with the PKA activator 6-Bnz-cAMP reduced GJ coupling. However, decreases in GJ coupling were prevented in islets treated with 6-Bnz-cAMP and S293 peptide (Figure 4H, p=0.056).

Taken together, these data demonstrate how the S293 peptide can restore declines in mouse and human islet GJ coupling and thus also suggest that Serine-293 on Cx36 may be a convergent target for both pro-inflammatory cytokines and the FFA palmitate to reduce GJ coupling.

### Inhibition of Cx36 Serine-293 maintains islet function and cell viability

We next tested whether using the S293 peptide to prevent GJ uncoupling in the context of a diabetogenic environment would preserve synchronized Ca^+2^ dynamics and β-cell viability, as occurs under genetic or pharmacological elevation in Cx36 GJ coupling (Figure 3). In addition to the effects on GJ coupling and [Ca^2+^], incubation of mouse islets with proinflammatory cytokines increases cell death and decreases GSIS (Figure S3). Incubation with the S293 peptide in the presence of cytokines restored [Ca^2+^] oscillatory dynamics (Figure 5A), and significantly increased the proportion of the islet that was active in response to glucose (Figure 5B, p=0.0009). In contrast, incubation with the scrambled peptide resulted in disorganized and unsynchronized [Ca^2+^] dynamics remaining (Figure 5A). Incubation with the S293 peptide in the presence of cytokines significantly reduced cytokine-induced β-cell death, by a factor of ∼40% (Figure 5C, p=0.0003). The reduction in β-cell death by incubation with the S293 peptide was not observed in islets from Cx36 knockout mice (Figure 5D). This is consistent with a role for Cx36 in protecting against apoptosis (Figure 3A, (Klee *et al*., 2011a)). Finally, incubation of mouse islets with cytokines and S293 peptide restored GSIS at 20 mM glucose (Figure 5E, p= 0.0003). In islets from Cx36 knockout mice incubation with the S293 peptide in the presence of cytokines led to a small elevation in GSIS, although this increase was not statistically significant (Figure 5F, p=0.11).

**Figure 5:**
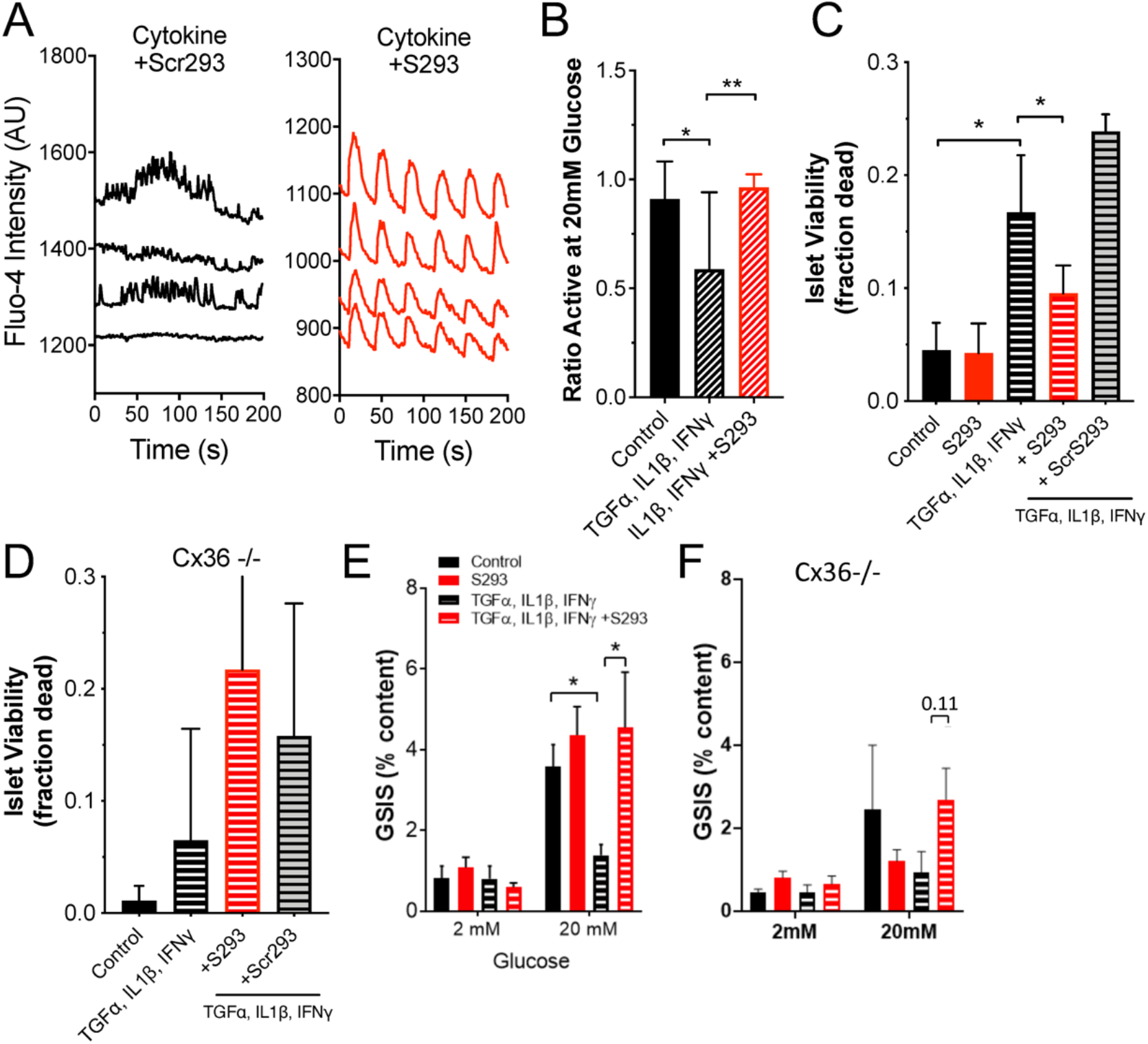
Inhibition of Cx36 Serine-293 maintains islet function and cell viability. A) Representative [Ca^2+^] time courses from four individual cells within an islet treated with a cocktail of 1 ng/ml mouse recombinant TNF-α, 0.5 ng/ml IL-1β, 10 ng/ml IFN-γ plus 1 μM scrambled peptide (cytokine+Scr293, left) or within an islet treated with a cocktail of 1 ng/ml mouse recombinant TNF-α, 0.5 ng/ml IL-1β, 10 ng/ml IFN-γ plus 1 μM S293 peptide (cytokine+S293, right). B) Area of [Ca^2+^] activity normalized to islet size (fraction active) for islets treated with TNF-α, IL-1β, = IFN-γ plus = S293 peptide, as in A. C) Islet viability, as indicated by the proportion of cells labelled as dead, under a cocktail of 10 ng/ml mouse recombinant TNF-α, 5 ng/ml IL-1β, 100 ng/ml IFN-γ and/or 1 μM S293 peptide or 1 μM scrambled (Scr) peptide. D) As in C for islets from Cx36 knockout mice. E) Glucose-stimulated Insulin secretion (GSIS) in islets treated with a cocktail of 1 ng/ml mouse recombinant TNF-α, 0.5 ng/ml IL-1β, 10 ng/ml IFN-γ plus 1 μM S293 peptide. F) As in E for islets from Cx36 knockout mice. Data in B representative of islets from n=5 mice; data in C representative of islets from n=6 mice; data in D representative of islets from n=8 mice; data in E representative of islets from n=5 mice; data in F representative of islets from n=6 mice. In B ** represents p=0.021; in C ** represents p=0.0003; In E ** represents p=0.0003; comparing groups indicated (all unpaired Student’s t-test). Data in B-F presented as mean±SD.

Together, these data support that hypothesis that preventing GJ uncoupling precludes the disruption to coordinated [Ca^2+^] dynamics, GSIS and islet cell death that is observed when islets are exposed to a diabetogenic environment.

### Inhibition of Cx36 Serine-293 maintains islet function in db/db mouse islets

The pathophysiology of T2D is complex and multifactorial (DeFronzo *et al*., 2015), thus culture of islets with pro-inflammatory cytokines or FFAs alone have a limited ability to model the islet microenvironment during T2D. We therefore tested the ability of S293 peptide to restore GJ coupling and [Ca^2+^] dynamics in islets from the db/db animal model of T2D. In islets isolated from 4-week old db/db mice the S293 peptide restored regular oscillatory [Ca^2+^] dynamics compared to in untreated db/db islets (Figure 6A). However, these oscillations had increased frequency compared to untreated control islets. When quantified, S293 peptide treatment in db/db islets increased the total fraction of the islet that was coordinated compared to untreated db/db islets (Figure 6B, p=0.002). Consistent with these data, we observed that S293 protected against decreases in GJ coupling in db/db islets. Significant decreases in GJ coupling were observed in untreated or scrambled peptide-treated db/db islets compared to equivalently treated control BLKS islets (Figure 6C; p=0.05 for BKS vs. db/db, p=0.067 for db/db vs. db/db+S293). However no decrease was observed in scrambled peptide-treated db/db islets compared to equivalently treated control BLKS islets. Thus in pre-diabetic db/db mouse islets, S293 peptide mimetic can restore both GJ coupling and [Ca^2+^] dynamics.

**Figure 6:**
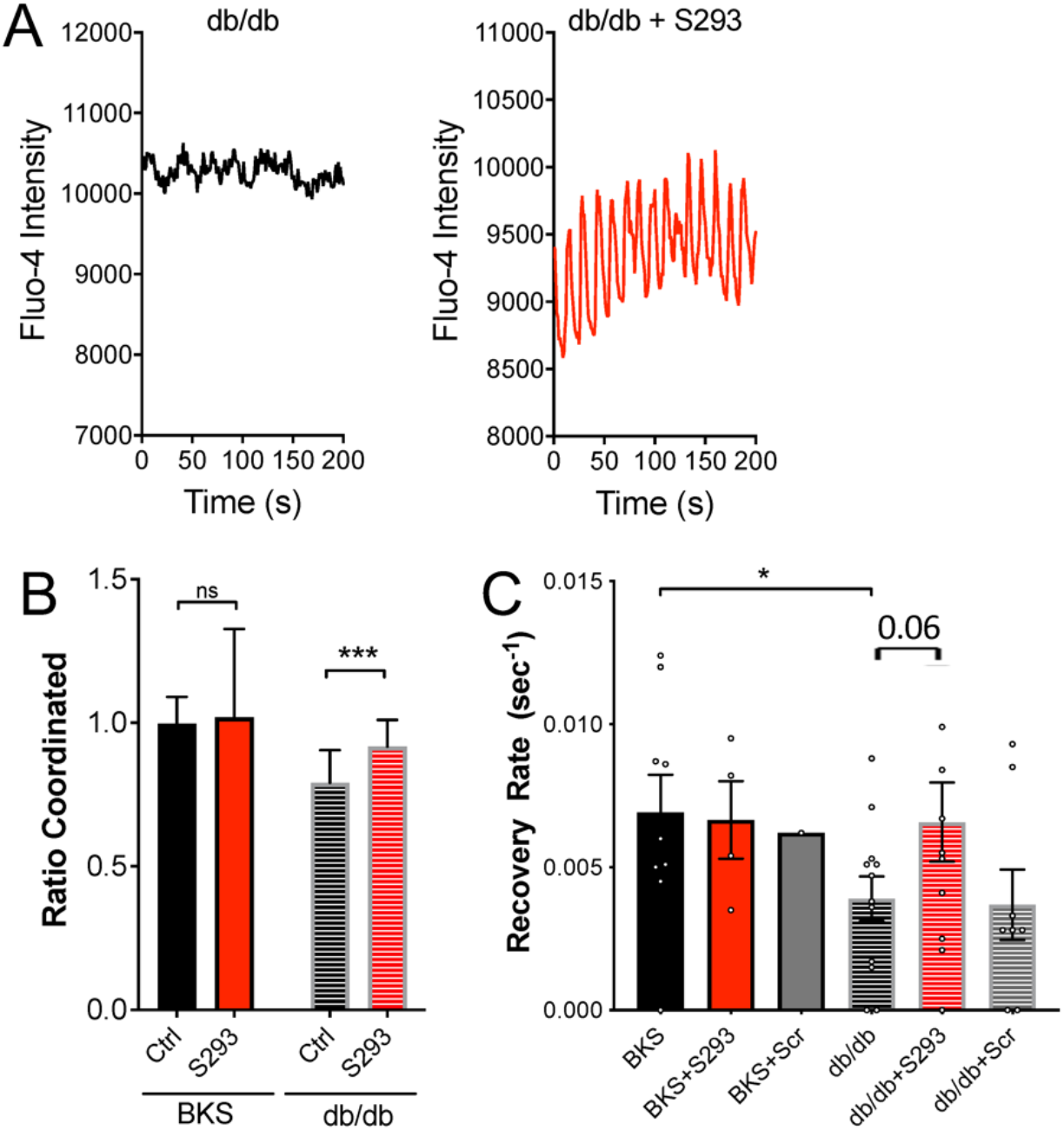
Inhibition of Cx36 Serine-293 maintains islet function in db/db mouse islets. A) Representative [Ca^2+^] time courses averaged over an islet from a 4w db/db mouse, that is either untreated (left) or treated with S293 peptide (right). B) Area of coordinated [Ca^2+^] normalized to islet size (ratio coordinated) for islets from 4w C57BLKS (WT) or db/db mice that are either untreated (Ctrl) or treated with S293 peptide as in A. Data is normalized to paired untreated C57BLKS islets. C) Mean gap junction permeability, as measured by FRAP recovery rate, for islets isolated from 4w C57BLKS mice (BKS) that are untreated, treated with the S293 peptide (+S293), or treated with the scrambled peptide (+Scr); and for islets isolated from 4w db/db mice that are untreated, treated with the S293 peptide (+S293), or treated with the scrambled peptide (+Scr). Data in B representative of islets from n=5 mice; data in C representative of islets from n=5 mice. In B *** represents p=0.002; in C * represents p=0.05; comparing groups indicated (all unpaired Student’s t-test). Data in C presented as mean±s.e.m. with each data point representing an islet. Data in B presented as mean±SD.

### Inhibition of Cx36 Serine-293 restores GJ coupling and GSIS in type2 diabetic human islets

We finally tested the ability of S293 peptide to restore the function of islets isolated from human donors with confirmed T2D. As described before we observed a small elevation in GJ coupling in islets from all healthy human donors upon incubation with S293 peptide (Figure 7A). Incubation of T2D human islets with S293 peptide restored GJ coupling with varying degrees (Figure 7B). No increase in GJ coupling was observed upon incubation of T2D human islets with scrambled peptide (Figure 7C). There was no significant change in glucose-stimulated insulin secretion in islets from healthy donor following S293 treatment (Figure 7D). The change in GSIS was highly variable following S293 treatment among the islets from T2D donors. However, when we separately analyzed GSIS data from islets isolated from the donors that showed a recovery in GJ coupling (an increase in recovery rate by >1.1), 3 out of 4 “responders” showed an increase in GSIS (Figure 7E). In contrast none of the “non-responders” showed an increase in GSIS (Figure 7E). There was also no significant change in glucose-stimulated insulin secretion in islets from T2D following scrambled peptide treatment (Figure 7F).

**Figure 7:**
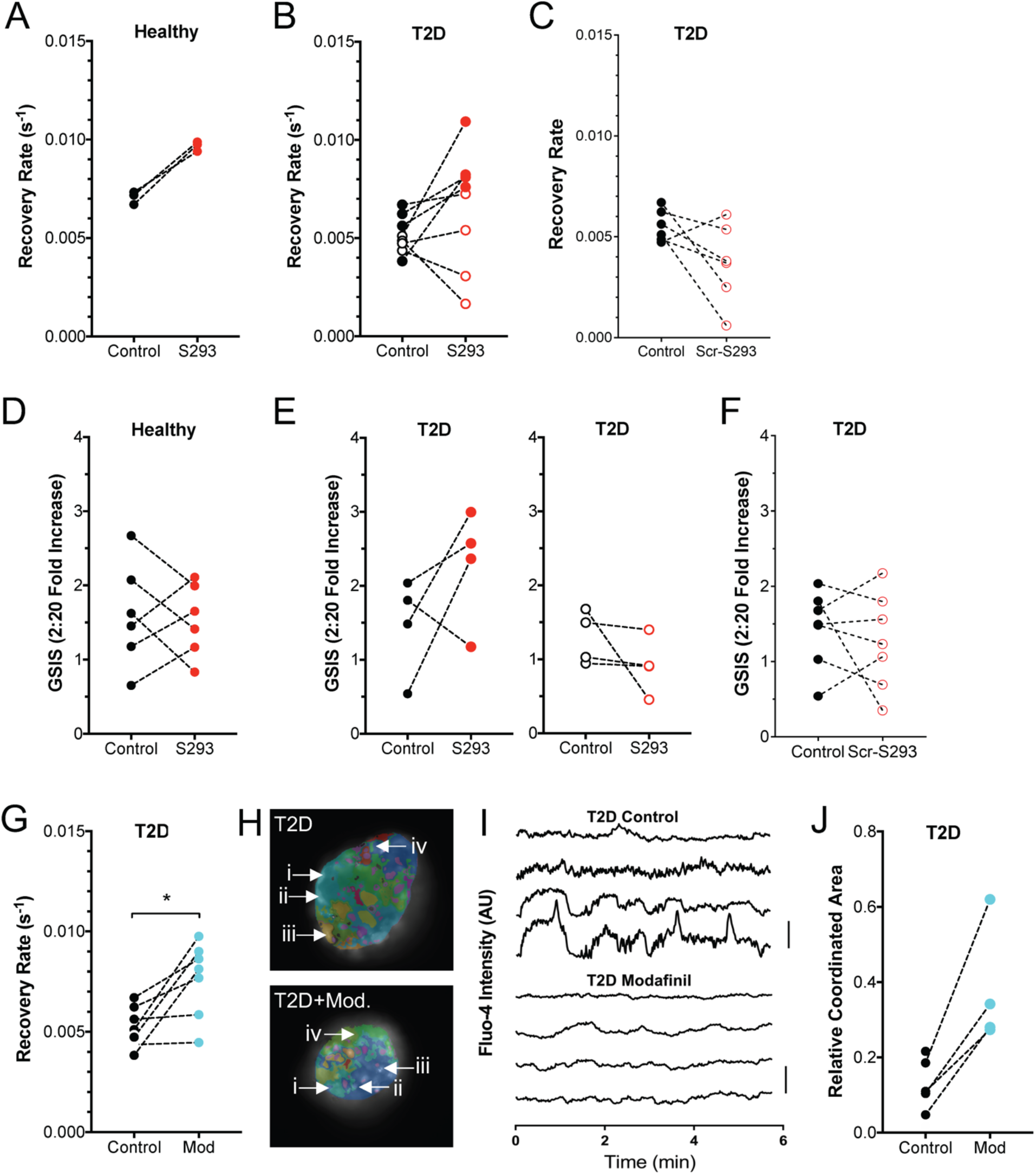
Inhibition of Cx36 Serine-293 restores GJ coupling and GSIS in type2 diabetic human islets. A) Gap junction permeability, as measured by FRAP recovery rate, for human islets isolated from healthy donors, that are either untreated (Control) or treated with S293 peptide. Data is presented as the mean recovery rate over islets from each donor, with dashed line connecting data for each donor. B) As in A for human islets isolated from donors with T2D, that are either untreated or treated with S293 peptide. C) As in A for human islets isolated from donors with T2D, that are either untreated or treated with scrambled peptide (Scr-S293). D) Glucose-stimulated Insulin secretion (GSIS) for human islets isolated from healthy donors, that are either untreated (Control) or treated with S293 peptide, as in A. E) GSIS as in D, for human islets isolated from T2D donors, that are either untreated (Control) or treated with S293 peptide. Data is presented as the mean recovery rate over islets from each donor, with dashed line connecting data for each donor. Left corresponds to those donors in B that showed a >10% elevation in GJ coupling by S293 peptide. Right corresponds to those donors in B that showed a decrease or <10% elevation in GJ coupling by S293 peptide. F) GSIS as in D, for human islets isolated from T2D donors, that are either untreated (Control) or treated with scrambled peptide (Scr-S293). G) Gap junction permeability for human islets isolated from donors with T2D that are untreated (control) or treated with 100μM modafinil (Mod). H) Representative false color map of [Ca^2+^] coordination in human islet from a T2D donor that is either untreated (T2D) or treated with 100μM modafinil (T2D+Mod). [Ca^2+^] activity is represented by presence of false color, with each color representing a separate region of [Ca^2+^] coordination, as in Figure 1 (see legend therein). I) Time courses from individual cells (i–iv) indicated in each of these islets. J) Largest area of coordinated [Ca^2+^] activity normalized to islet size (relative coordinated area) averaged over human islets from each T2D donor that are untreated (control) or treated with 100μM modafinil (Mod). Data in A representative of 3 donors; data in B representative of 8 donors; data in C representative of 6 donors; data in D representative of 6 donors; data in E representative of 4 donors each; data in F representative of 7 donors; data in G representative of 7 donors; data in J representative of 4 donors. In G * represents p=0.018; in J * represents p=0.04; comparing groups indicated (paired Student’s t-test). Data in A-G, J presented as one point per donor.

We also tested whether modafil would provide similar recoveries in islet function. We observed a highly variable increase in GJ coupling in islets from T2D donors following treatment with modafinil (Figure 7G). Following [Ca^2+^] imaging, we also observed a substantial increase in the fraction of the islet that was coordinated in islets from T2D donors (Figure 7H).

These data show that the S293 peptide mimetic, as well as pharmacological elevations in GJ coupling were able to restore [Ca^2+^] dynamics and glucose-stimulated insulin secretion in islets from human donors with T2D.

## DISCUSSION

Type2 diabetes results from insufficient secretion of insulin in the context of insulin resistance, followed by subsequent further decline in β-cell mass. Dysfunction to β-cells within the islet of Langerhans is one main cause for this lack of sufficient insulin production. Cx36 gap junction (GJ)-mediated electrical coupling is critical to the regulation of pulsatile insulin secretion. GJ coupling is diminished by conditions associated with diabetes, including by inflammatory cytokines, lipotoxic conditions, in animal models of diabetes and in aged human islets (Hodson *et al*., 2013; Farnsworth *et al*., 2016a; Westacott *et al*., 2017; Corezola do Amaral *et al*., 2020). In this study we sought to further define how conditions associated with diabetes disrupt GJ coupling within the islet and determine how a disruption to GJ coupling impacts downstream [Ca^2+^], insulin secretion and cell viability under diabetogenic conditions. We found that transgenic over-expression of Cx36, pharmacological activation of GJ coupling, and a novel peptide mimetic that restores Cx36 GJ coupling all protect against islet dysfunction by retaining [Ca^2+^] dynamics and cell viability.

### Importance of GJ coupling to islet function in health and in diabetes

Gap junction channels are critical for coordinating and enhancing glucose-regulated electrical activity and pulsatile insulin secretion across the islet. Cx36 deficient mice lack any electrical coupling between β-cells in the islet, highlighting how Cx36 gap junctions are the driver of this electrical coupling. These mice are glucose intolerant as a result of both diminished first phase insulin secretion and disrupted pulsatile insulin secretion (Head *et al*., 2012). Thus, diminished Cx36 GJ coupling under diabetogenic conditions, and in subjects with T2D, would disrupt pulsatile insulin secretion and glucose tolerance. Indeed loss of first phase secretion is a hall mark of pre-T2D, and individuals with T2D show diminished pulsatile insulin as in Cx36 null animals (Menge *et al*., 2011). Thus, loss of Cx36 GJ coupling likely explains, in part, the reduced glucose tolerance in T2D,

Cx36 GJ coupling is also protective against cell death induced by cytotoxic conditions such as by proinflammatory cytokines (Klee *et al*., 2011a). Indeed, we demonstrated through multiple means that elevating Cx36 GJ coupling or protecting against its decline reduces apoptosis (Figure 3, 5). Thus, the loss of GJ coupling in diabetes likely exacerbates cell death and islet decline. Interestingly however, we did not find a strong role for Cx36 GJ coupling to be protective against declines in the regulation of insulin secretion induced by proinflammatory cytokines (Figures 3,5), but did see a role for Cx36 GJ coupling to be protective against disrupted insulin secretion in T2D subjects (Figure 7). Thus the loss of GJ coupling in diabetes may also influence the regulation of insulin release.

Through utilizing multiple approaches and models of diabetes we can have high confidence as to the protective role provided by Cx36 GJ coupling. We observed improvements in cell viability and GSIS using the Cx36:S293 mimetic, which were not observed in Cx36 deficient islets. Data from Cx36 deficient islets indicates that potential off-target effects of the Cx36:S293 mimetic are unlikely to contribute significantly to cell death, but may contribute slightly to GSIS. Qualitatively similar results in terms of cell death, GSIS and [Ca^2+^] following Cx36 over expression and following modafinil-based increases in GJ coupling (which may have off target effects) support Cx36 playing a key role.

It is important to note that we solely utilize in vitro based experiment’s in this study, combining mouse and human islet models. However, there are challenges to examining our questions invivo. Firstly, the Cx36 over-expression decreased slightly under inflammatory conditions and thus invivo protection against declines in GJ coupling may be limited. Peptides have low half-life in circulation and thus the S293 mimetic would likely provide limited action following systemic delivery. Modafinil would also have other actions invivo, as it can impacts CNS function. As such whether the protection against apoptosis and disruptions to GSIS is sufficient to impact diabetes is still to be determined. As a second point, while Cx36 is the predominant connexin in both mouse and human beta-cells (Miranda *et al*., 2022), the precise role for Cx36 GJ coupling in human islets is not fully determined. Other mechanism have also been suggested to contribute to synchronizing [Ca^2+^] dynamics in the human islet via paracrine communication (Rodriguez-Diaz *et al*., 2011; van der Meulen *et al*., 2015; Almaça *et al*., 2016; Menegaz *et al*., 2019). We have previously demonstrated that improving GJ coupling using pharmacological activators in aged human islets improves [Ca^2+^] regulation and first phase GSIS (Westacott *et al*., 2017). Nevertheless, we still observed significant variability among human donors, with batches responding effectively to the S293 mimetic and/or modafinil (responders) and batches not (non-responders). This response did not significantly dependent on the state of the islets as measured by GSIS or [Ca^2+^], but may depend on some other factor.

Overall, increasing Cx36 GJ coupling or protecting against its decline protects against disruptions to islet [Ca^2+^], GSIS and cell viability, across both mouse and human islets.

### Regulation of Cx36 by S293 peptide mimetic

We demonstrated that treatments that restore or elevate Cx36 GJ coupling are protective against declines in islet function ([Ca^2+^], GSIS in human T2D) and cell death. Key points worthy of discussion are how mechanistically our treatments elevate GJ coupling and how elevated GJ coupling provides such protection.

Robust protection is provided by the S293 mimetic peptide. This mimetic peptide was designed to provide a competitive substrate for kinases that would normally phosphorylate the native S293 site, as well as other regulators of Cx36 that would bind this region. This S293 site has been identified as a critical for the regulation of Cx36 GJ coupling in AII amacrine cells, where it is phosphorylated by PKA (Urschel *et al*., 2006). It is also identified as a phosphorylation site for PKCδ. PKA is activated within human islets during lipotoxic conditions and mediates the disruption to GJ coupling (Hodson *et al*., 2013). We show that the S293 peptide can block the decrease to GJ coupling induced by PKA activation (Figure 4). Similarly, PKCδ is activated within mouse and human islets during inflammatory conditions and mediates the disruption to GJ coupling (Farnsworth *et al*., 2016a). Therefore, the S293 peptide mimetic may be competing out factors that either bind or phosphorylate the key regulatory site(s) on the Cx36 cytoplasmic tail that regulates channel closure or trafficking. The mechanisms by which modafinil promotes electrical coupling are less clear. Reports in studying its action on Cx36 in the CNS suggest activation of CamKII and phosphorylation sites distinct from those phosphorylated by PKA/PKC. Thus, phosphorylation sites that promote Cx36 channel opening may be targeted. However, microtubule binding to this c-terminal region has been reported (Brown *et al*., 2019), and thus regulation of trafficking is also possible. Indeed, the S293 peptide blunts the decrease in membrane localized Cx36 induced by pro-inflammatory cytokines, which is consistent with improved trafficking of Cx36. Detailed phospho-proteomic measurements will be required in future studies to delineate whether the S293 site or other sites are phosphorylated or dephosphorylated by diabetogenic conditions (pro-inflammatory cytokines, FFA, db/db mice, T2D donor) and the interventions we make (Cx36:S293 peptide, modafinil). We also note that a Cx43 peptide mimetic (alphaCT1) can promote gap junction aggregation (Hunter *et al*., 2005); and we cannot exclude that the S293 peptide also directly regulates membrane-localized Cx36 organization. However the limited impact of the S293 peptide under baseline conditions (Figure 4) argues against this.

The other major question is how Cx36 increases providing protection against cell death. Retaining normal GJ coupling would maintain normal coordinated [Ca^2+^] dynamics, and this maintained [Ca^2+^] would contribute to improved GSIS. Cx36 GJ coupling also protects against apoptosis, as we demonstrate here and we and others have previously demonstrated (Klee *et al*., 2011a). The mechanism underlying this are poorly defined. However, published studies from our lab suggest the increased cytosolic [Ca^2+^] at lower glucose levels induced by loss of Cx36 increases the susceptibility to apoptosis (Farnsworth *et al*., 2022). This is consistent with well-established results in which elevated cytosolic [Ca^2+^] and calcineurin activation contributes to intrinsic apoptotic mechanisms (Wang *et al*., 1999).

### Relevance to diabetes and greater context

The decline in functional β-cell mass is important in the pathophysiology of diabetes, determining overt disease onset, progression and success of treatment. Therapies are needed to preserve β-cell mass and existing β-cell function: such therapies would prolong the duration until diabetes onset and thus lower the risk of diabetic complications, as well as lessening the financial and physical burden to the patient. Our results suggest that targeting Cx36 GJ coupling, and particularly targeting it to prevent its disruption in T2D can preserve normal GSIS dynamics and cell viability.

Targeting Cx36 is also strongly supported by its role in the pathogenesis in T2D. Several studies, including results presented here, demonstrate its disruption under diabetogenic conditions (Carvalho *et al*., 2012; Hodson *et al*., 2013; Farnsworth *et al*., 2016a; Westacott *et al*., 2017; Corezola do Amaral *et al*., 2020). We further demonstrate its disruption in human donors with T2D compared to age matched healthy donors (Figure 1). Our results also suggest this is linked to the development of T2D and not as a result of subsequent poor glucose control given lack of any correlation with HbA1c. Importantly the improvement in function after targeting Cx36 via the mimetic peptide or via modafinil further support an important role in human T2D. In addition to these results, gene variants in the *GJD2* gene (encoding Cx36) are associated with T2D (Cigliola *et al*., 2016). The disease associated variant has reduced coupling, further supporting a role for Cx36 in T2D and thus targeting it for T2D prevention.

The most effective strategy we found to target Cx36 GJ coupling was the Cx36:S293 mimetic peptide. This showed effective protection under diabetogenic conditions in both mouse and human islets, including from T2D donors. However, we were unable to effectively test it invivo given the short circulation time of peptides before clearance. Delivering peptides for therapeutic use is a major challenge: once-weekly delivery systems such as for GLP1R agonists are complex. Implementing such a delivery system, designing a more stable mimetic peptide, or targeting the peptide directly to the islet microenvironment will be needed in future work. Modafinil which is an approved drug, also provided protection in a range of exvivo models and assessing its effectiveness in human patients at risk for T2D would be useful. Our results examining transgenic over-expression of Cx36 also suggest a potential challenge in using this method to assess the role of Cx36 in protecting against T2D progression invivo. While protection against apoptosis was provided by transgenic over-expression of Cx36, GJ coupling still did decrease significantly under diabetogenic conditions. We did not test whether this over-expression could still protect against islet decline in animal models of T2D. However, we have observed a minor protection against disease protection in animal models of T1D (Farnsworth *et al*., 2022). In this case autoimmunity is still progressing and other means to target β-cell death do not completely abrogate disease progression.

A general limitation in treating T2D is the fact that the disease is complex and multifactorial, with no single pathway dominant in regulating islet decline. Thus, targeting solely Cx36 GJ coupling is unlikely to be sufficient to fully abrogate disease. Furthermore, even if we can preserve beta cell function, the continuing metabolic stress caused by insulin resistance will likely mean that the preservation will be temporary. Nevertheless, targeting Cx36 GJ coupling could be effective as part of a broader strategy, such as targeting ER stress and stimulating GSIS.

## ACKNOWLEDGEMENTS

The authors acknowledge the guidance and expertise of Dr. Robert Hodges, Director of the University of Colorado Protein Peptide Synthesis core for his guidance in the design and synthesis of both S293 and scrambled mimetic peptide sequences.

## AUTHOR CONTRIBUTIONS

JRS designed and performed experiments, analyzed the data, wrote the manuscript; MJW performed experiments, analyzed data; JM performed experiments, analyzed data; NLF performed experiments, analyzed data; VK performed experiments, analyzed data; WES performed experiments; CHL performed experiments, analyzed data; JMD analyzed data; AH performed experiments; NL analyzed data; RKPB designed experiments, wrote the manuscript.

## DATA AVAILABILITY

All raw data will be made available upon request.

---

**Abstract figure:**
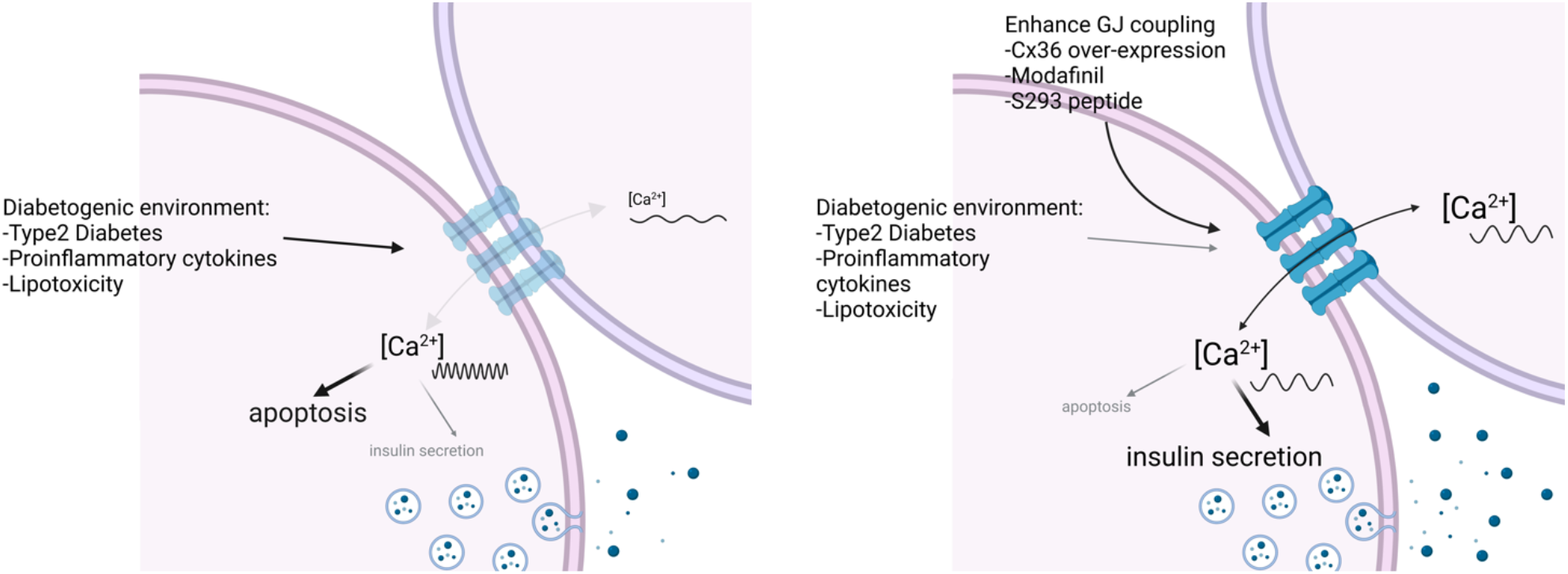
Left: Diabetogenic environments disrupt Cx36 gap junction coupling, Ca^2+^ activity/coordination and insulin secretion while increasing apoptosis. Right: Elevating Cx36 via genetic means, Modafinil or S293 peptide recovers Ca^2+^ activity/coordination and insulin secretion while reducing apoptosis.

## SUPPLEMENTAL MATERIALS

**Supplemental Table 1:** List of human donors from which islets were examined in this study, with the experiments performed and treatments applied and whether they responded to treatment.

| Donor ID | UNOS ID | Diabetes status | Experiments performed: |  |  | Treatments: |  |
| --- | --- | --- | --- | --- | --- | --- | --- |
|  |  |  | GSIS | Ca2+ | GJ FRAP | S293 pep | Modafinil |
| 993 | AA2482 | T2D | x | x | x |  |  |
| 1049 | ABDG032 | T2D | x | x | x |  |  |
| 1085 | ABFG183 | T2D | x | x | x |  |  |
| 1431 | ACJV399 | T2D | x | x | x | x (resp.) | x |
| 1562 | ADCD391 | T2D | x | x | x | x (resp.) | x |
| 1565 | ADCI176 | T2D | x | x | x | x (non.resp.) | x |
| 1576 | ADCY327 | T2D | x | x | x | x (resp.) | x |
| 1667 | ADGD159 | T2D | x |  |  | x (non.resp.) |  |
| 1808 | AECB422 | T2D | x | x | x | x (resp.) | x |
| 1816 | AEC2011 | T2D | x | x | x | x (non.resp.) | x |
| 1832 | AEDZ307 | T2D | x | x | x | x (non.resp.) | x |
| 971 | AAIQ245 | ND | x | x |  |  |  |
| 1060 | ABD1375 | ND | x | x |  |  |  |
| 1070 | ABEI419 | ND | x | x |  |  |  |
| 1085 | ABFG183 | ND |  |  | x |  |  |
| 1089 | ABFE184B | ND | x | x |  |  |  |
| 1099 | ABFV041 | ND | x | x |  |  |  |
| 1107 | ABGK387A | ND | x | x | x |  |  |
| 1118 | ABG2361 | ND | x | x |  |  |  |
| 1122 | ABHC283 | ND | x | x |  |  |  |
| 1162 | ABJR111 | ND |  |  | x |  |  |
| 1174 | ABJ5204 | ND | x | x |  |  |  |
| 1251 | ACBO251 | ND | x | x |  |  |  |
| 1271 | ACCR206A | ND | x | x |  |  |  |
| 1281 | ACCZ181-A | ND |  |  | x |  |  |
| 1310 | ACDH493 | ND | x | x |  |  |  |
| 1339 | ACEY057A | ND | x | x |  |  |  |
| 1357 | ACFO053-AA | ND |  |  | x |  |  |
| 1374 | ACGK380 | ND |  | x |  |  |  |
| 1401 | ACH4140 | ND | x | x |  |  |  |
| 1410 | ACIK323A | ND |  |  | x |  |  |
| 1503 | ACLE269 | ND | x | x |  |  |  |
| 1530 | ADAM295 | ND |  | x |  |  |  |
| 1536 | ADAU462B | ND | x | x |  |  |  |
| 1550 | ADBQ038 | ND | x | x |  |  |  |
| 1602 | ADDQ143 | ND |  |  | x |  |  |
| 1614 | ADEJ409 | ND |  | x |  |  |  |

**Supplemental Figure 1:**
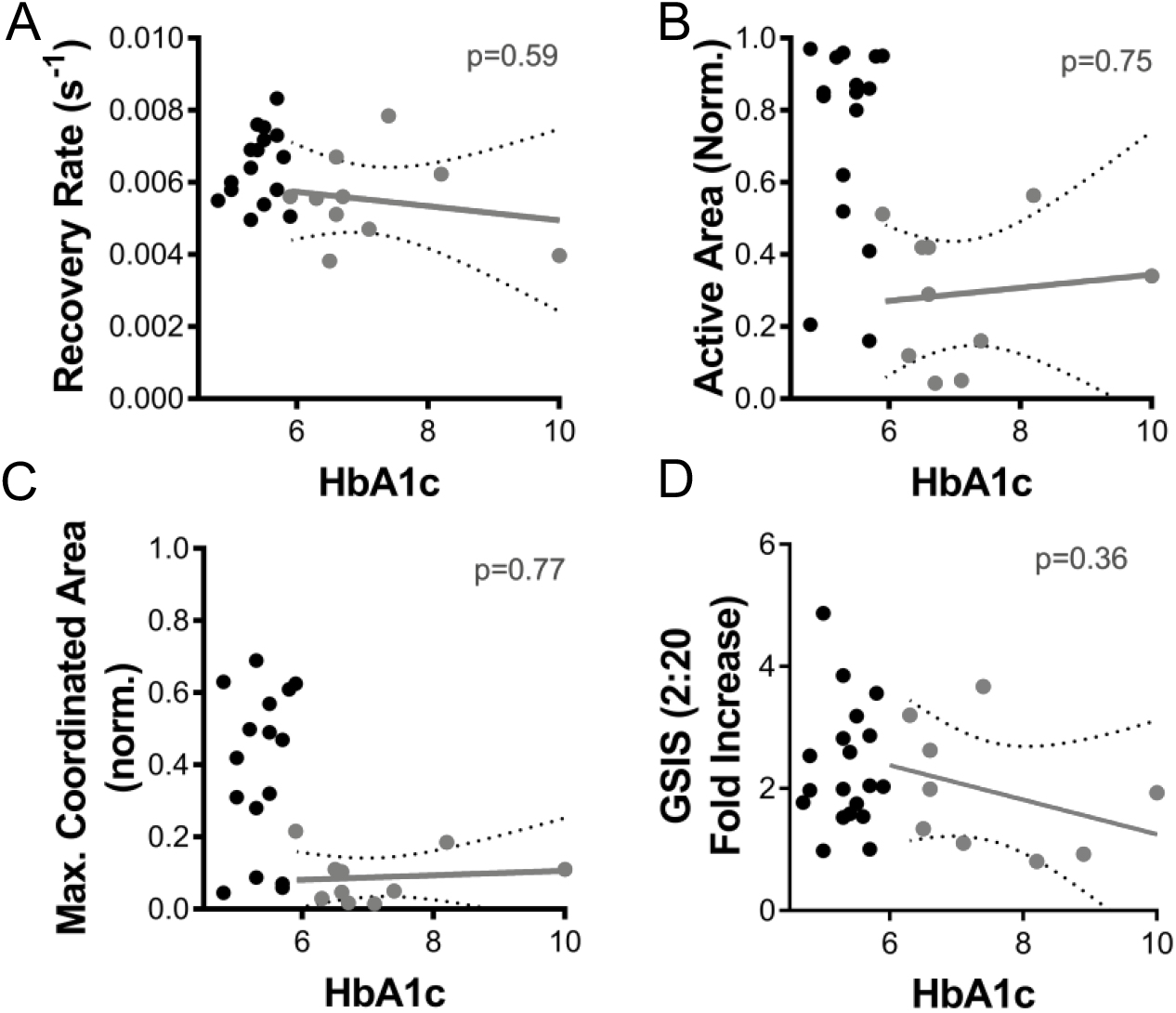
Link between donor measurements and HbA1c. A) Gap junction permeability, as measured by FRAP recovery rate, for human islets isolated from healthy donors (black) and T2D donors (grey); where HbA1c information is available. Regression (solid line) and 95% CI (dashed line) is calculated among T2D donors. B) As in A for area of [Ca^2+^] activity normalized to islet size (Active Area). C) As in A for largest area of coordinated [Ca^2+^] activity normalized to islet size (max. coordinated area). D) As in A for glucose-stimulated insulin secretion (fold change from 2mM to 20mM glucose). Data in A-C representative of 10 T2D donors and 16 healthy donors; data in D representative of 9 T2D donors and 19 healthy donors. p values describe significance of correlation between measurement and HbA1c for islets from T2D donors (F test).

**Supplemental Figure 2:**
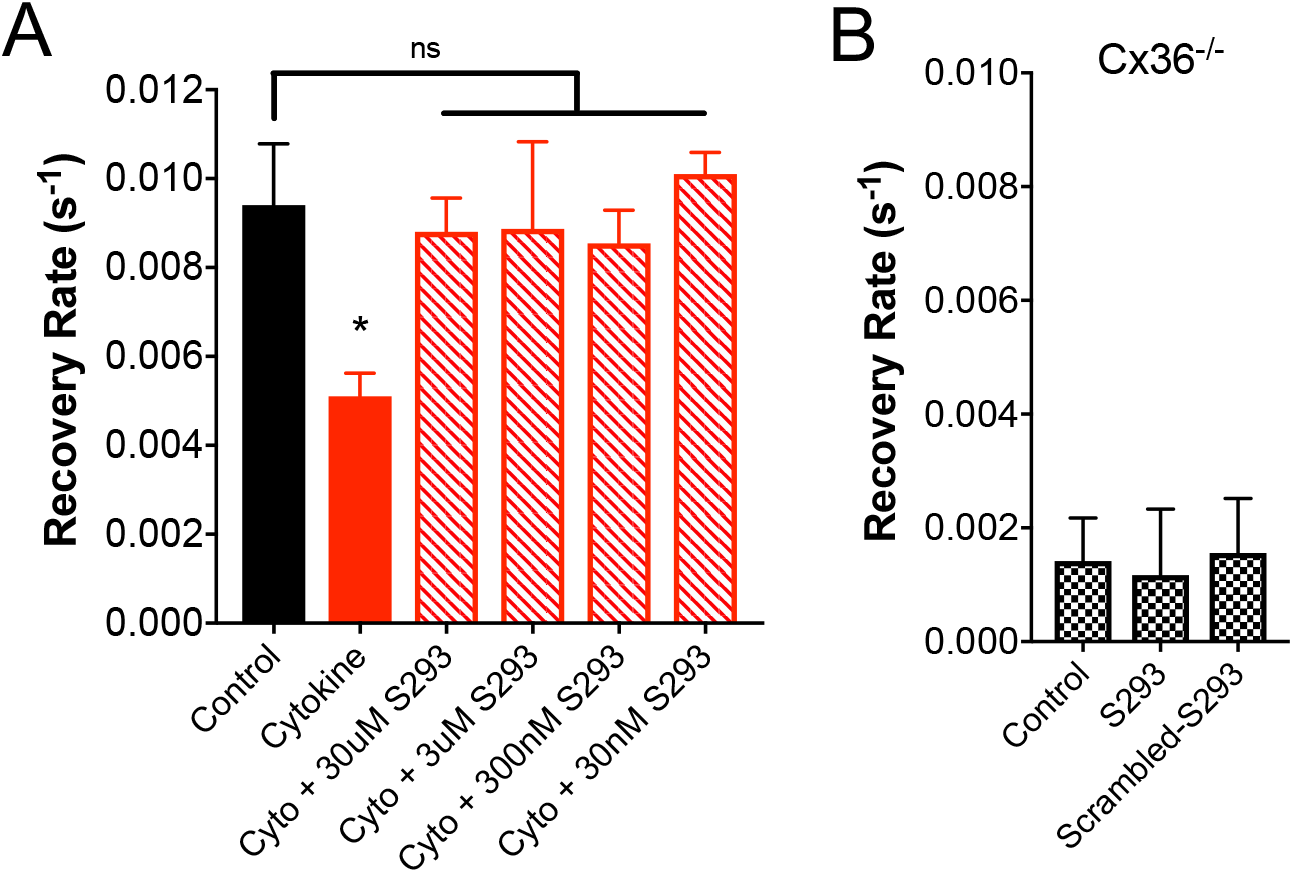
Dose response of S293 peptide. A) Gap junction permeability, as measured by FRAP recovery rate, for mouse islets that are untreated (control), treated with a cocktail of 1 ng/ml mouse recombinant TNF-α, 0.5 ng/ml IL-1β, 10 ng/ml IFN-γ, alone (Cytokine) or together with varying concentrations of the S293 peptide (Cyto+ S293, concentration as indicated). B) Gap junction permeability in islets from Cx36 knockout mice that are untreated, treated with S293 peptide or treated with scrambled peptide. Data in A representative of 4 islets per condition; data in B representative of 3 (S293) or 5 (scrambled) islets. * in A represents p=0.046; comparing groups indicated (unpaired Student’s t-test).

**Supplemental Figure 3:**
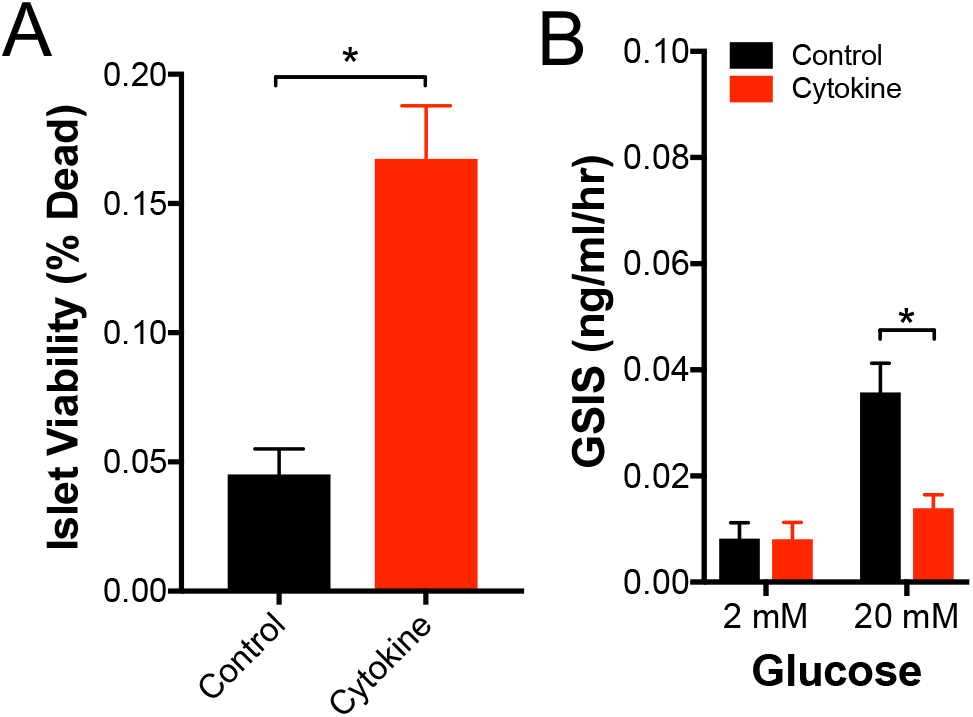
Additional characterization of cytokine effects on the islet. A) Islet viability, as indicated by the proportion of cells labelled as dead, for mouse islets that are either untreated or treated with a cocktail of 10 ng/ml mouse recombinant TNF-α, 5 ng/ml IL-1β, 100 ng/ml IFN-γ (Cytokine). B) Glucose-stimulated Insulin secretion (GSIS) for mouse islets that are treated as in B. Data in A representative of 6 mice; data in B representative of 5 mice. * in A represents p=0.003; * in B represents p=0.016 comparing groups indicated (unpaired Student’s t-test).

